# Sex Chromosome Turnover and Structural Interspecific Genome Divergence Shapes Meiotic Outcomes in Hybridizing *Cobitis*

**DOI:** 10.1101/2025.04.01.646337

**Authors:** S. A. Schlebusch, V. Trifonov, Z. Halenková, M. Klianitskaya, D. Dedukh, A. Ruiz Herrera, L. Álvarez González, G. Pujol Infantes, E. Hřibová, L. Andjel, O. Bartoš, P. Pajer, T. Tichopád, D. Kulik, J. Kotusz, M. Kaštánková Doležálková, A. Bohne, A. Marta, P. Horna, R. Reifová, Y. Guiguen, J. Pačes, K. Janko

## Abstract

It has been empirically established that genome mixing between divergent species can trigger meiotic aberrations, ultimately leading to the emergence of asexual reproduction through the production of unreduced gametes in various metazoan lineages. Yet, it remains poorly understood how such asexual hybrids cope with co-inherited differences in sex determination systems, diverged regulatory networks, and chromosomal incompatibilities— especially in the context of increased ploidy. Addressing these questions requires high-quality, chromosome-level reference genomes of the parental species involved in hybrid formation.

Here, we present the first chromosome-level genome assemblies for three hybridizing *Cobitis* species (*C. elongatoides*, *C. taenia*, and *C. tanaitica*), providing a comprehensive framework to investigate the genetic and cytogenetic basis of hybrid sterility and the transition to asexuality. By integrating genome scaffolding, male/female pooled sequencing, and molecular cytogenetics, we uncover extensive structural variation among homologous chromosomes of the three species, despite their overall syntenic conservation.

Population-level Pool-Seq analyses further revealed that each species possesses a distinct, non-homologous sex chromosome, highlighting sex chromosome turnover even among recently diverged lineages. These assemblies enabled the design of chromosome-specific painting probes, which we applied to meiotic metaphase I spreads of diploid hybrids. This approach revealed striking differences in the pairing success of orthologous chromosomes, with some (e.g., Ch01B) frequently forming bivalents, while others (e.g., Ch01A, Ch05, Ch20) failed to do so and remained unpaired.

Our results demonstrate that chromosome-specific features, shaped by structural evolution and sex-linked divergence, contribute unequally to hybrid meiotic failure. Together, this work provides a high-resolution genomic and cytogenetic framework to understand how interspecific hybridization gives rise to clonality, and how the architecture of inherited parental genomes shapes the success or breakdown of meiosis in hybrid vertebrates.

## Introduction

Reproduction, the ability to transmit a genome from one generation to another, is fundamental to all living organisms. In metazoans, this primarily occurs through the production of reduced gametes via meiotic divisions with recombination. This process likely evolved as an efficient means to repair DNA, but it also provides significant advantages over non-recombinant modes, such as generating variability to escape fast-evolving pathogens, removing deleterious mutations, and facilitating genetic exchange within a species’ gene pool (Lenormand et al., 2016). However, reproductive modes vary widely, and even meiosis and recombination frequencies are optimized for specific genomic regions, environments, or sexes (Kochakpour & Moens, 2008; Lenormand et al., 2016; Ortiz-Barrientos et al., 2016; Thompson & Jiggins, 2014). Many typically sexual species, including humans, transmit parts of their genome, like mitochondria and sex chromosomes, in an asexual manner with little or no recombination. In extreme cases, some multicellular organisms have abandoned sex altogether and instead produce unreduced gametes.

Hence, unlike sexually reproducing species, which likely share the fundamentals of sexual reproduction from a common ancestor (Bernstein & Bernstein, 2010), ‘asexual’ lineages do not represent a simply definable group. They are scattered across the tree of life and employ a wide spectrum of independently arisen cytological mechanisms for gamete production, ranging from completely ameiotic processes (apomixis) to those involving altered versions of meiotic divisions (automixis) (Stenberg & Saura, 2009, 2013), with different genetic consequences for the evolution and ecology of these lineages. Certain types of automixis involve homogenizing processes such as intragenomic recombination or exclusion of large portions of the genome, reducing heterozygosity in the asexual progeny (Bi & Bogart, 2006). In contrast, automixis involving premeiotic endoreplication (PMER) results in progeny genetically identical to the mother, as genetic material is duplicated before meiosis and recombination occurs between sister (Arai & Fujimoto, 2013; Dedukh et al., 2020; Lutes et al., 2010).

Despite the great variability of asexual organisms and their polyphyletic origins, recent research has identified several emerging trends, with unrelated asexual organisms often sharing similar cytogenetic and molecular traits. For example, the genomes of asexual organisms often do not accumulate deleterious mutations as quickly as originally expected (Jaron et al., 2021; Kočí et al., 2020; Loewe & Lamatsch, 2008; Pellino et al., 2013). Another such trend is that the abandonment of sex frequently coincides with interspecific hybridization, which, for reasons yet unknown, is correlated with increasing divergence between hybridizing sexual species (Marta et al., 2023; Moritz et al., 1989). For many such hybrids, “asexual” production of unreduced gametes is achieved by the same cytological mechanism, the PMER (Arai & Fujimoto, 2013; Dedukh et al., 2020; Lutes et al., 2010). Interestingly, this process is generally sex-specific and typically confined to females, while hybrid males from the same crosses are usually unable to produce clonal gametes (Stöck et al., 2021; Tichopád et al., 2022).

The switch to asexuality may also be accompanied by changes in genome and gene expression which are consistent across independently arisen lineages (Parker et al., 2019). Once asexuality appears in a lineage, the genomes originally inherited from sexual ancestors do not remain static but rather tend to accumulate structural changes. Counter intuitively, this can often lead to a loss of heterozygosity due to processes such as gene conversion (Janko et al., 2021; Jaron et al., 2021; Warren et al., 2018). Comparisons with sexual ancestor species have shown that gene conversions in asexual genomes are tightly correlated with the expression of orthologous alleles, base composition of loci, and specific gene functions (Janko et al., 2021). This adaptive loss of heterozygosity may selectively remove certain parental alleles, leading to the balancing and optimization of components in regulatory networks in hybrid genomes (Bartoš et al., 2019). Understanding genome evolution under restricted recombination, especially in asexuals, represents a dynamic research field that necessitates proper comparative contrasts between asexuals and their direct sexual ancestors. Despite advances in whole genome sequencing, many lineages still lack properly assembled and annotated genomes, let alone characterization of structural variants and distribution of repetitive elements. This challenge is further complicated by the fact that many asexuals used in comparative studies either lack direct sexual counterparts or arose as interspecific hybrids combining genomes from distinct sexual species. Without being able to compare genotypes and phenotypes of contemporary asexual strains to their sexual progenitor populations, it is difficult to discern whether observed commonalities among unrelated asexual lineages are coincidental or indicative of deeply shared mechanisms enabling parallel switches to asexuality in distant organisms (Albertini et al., 2019; Janko et al., 2018; Murphy et al., 2000; Stöck et al., 2021). Acquiring high-quality reference genomes of carefully determined and selected parental species and comparing them to asexuals is essential for addressing these questions.

The spined loaches of the genus *Cobitis* serve as an appealing model for understanding the link between speciation, hybridization, sex, and asexuality, with several instances of interspecific hybridization found across Eurasian hybrid zones, between species that diverged between 1 and 15 million years ago (Figure 1). This hybridization produces either fertile sexual hybrids among closely related species or, in cases of greater divergence, sterile hybrid males and fertile but asexually reproducing hybrid females. These asexual females are found in diploid, triploid, and tetraploid states and all produce gametes via PMER to produce clonal eggs. Hybridization leading to asexuality has been ongoing throughout the Pleistocene, resulting in many clonal strains, some of which originated recently while others are several hundred thousand generations old (Janko et al., 2012). While clonal hybrids tend to conserve their inherited parental karyotype structure for thousands of generations without significant restructuring (Majtánová et al., 2016), they are subject to a gradual loss of heterozygosity, which accumulates selectively in certain genic groups and depends on the relative transcription of the alleles (Janko et al., 2021).

**Figure 1.**
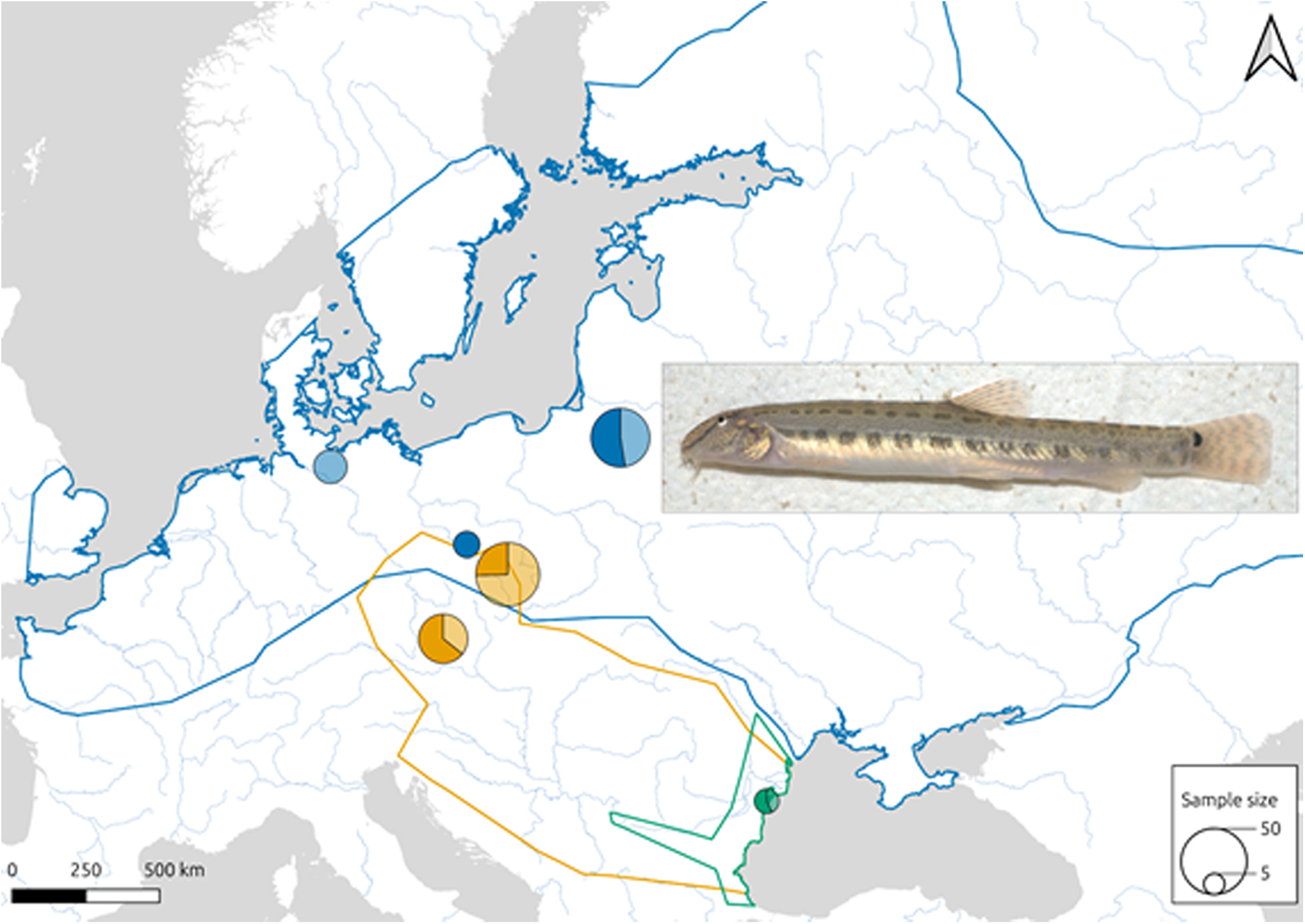
Map of European rivers indicating the distribution ranges of the three *Cobitis* species included in this study. *C. taenia* is in blue, *C. tanaitica* is in green and *C. elongatoides* is in yellow. Pie charts indicate the sample size and sex ratio of samples taken from each locality (males are indicated by the darker colour and females by the lighter one). The insert indicates a *C. taenia* female individual.

The sex determination system in these hybrids remains unknown, but clonal gametogenesis appears strictly sex-biased: only hybrid females undergo PMER, while their hybrid brothers fail to do so and arrest gametogenesis due to parental chromosome incompatibility (Dedukh et al., 2020; Marta et al., 2023). Interestingly, spermatogonial germ cell transplantation from sterile hybrid males into female gonads enables them to undergo PMER within the female gonadal environment (Tichopád et al., 2022).

These patterns raise important questions about the extent to which genome evolution in asexuals is affected in general by heredity without recombination versus being driven by the properties and divergence of genomes inherited from sexual ancestors, including specific genetic sex determination mechanisms, structural variants affecting chromosome pairing, divergence in repetitive elements, gene regulatory networks, and epigenetic signalling (Janko et al., 2024; Tichopád et al., 2022).

The aim of this study is to generate high-quality, chromosome-level genome assemblies for three parental *Cobitis* species—*C. elongatoides*, *C. taenia*, and *C. tanaitica*— which serve as the sexual progenitors of various asexual hybrid lineages. These assemblies provide a necessary foundation for investigating the genomic basis of hybrid sterility and asexuality, focusing on four key aspects: the extent and nature of structural variants (SVs) accumulated between species and their impact on chromosomal compatibility; divergence in repeat element content and dynamics across lineages; identification and comparative analysis of genetic sex determination systems; and chromosomal pairing behavior in hybrid males as a mechanistic insight into meiotic failure.

In addition to addressing these biological questions, our work fills a major taxonomic and geographic gap in genome availability for the Cobitidae family. Despite its species richness and unique reproductive biology, high-quality chromosome level genomic assemblies for Cobitidae remain scarce and restricted to two East Asian representatives - *Paramisgurnus dabryanus* and *Misgurnus anguilicaudatus* (Sun et al., 2024; L. Zhang et al., 2025). By providing the first chromosome-level assemblies from the western Palearctic *Cobitis* lineage, our study more considerably increases the number of Cobitidae reference genomes and opens the door to broader comparative and evolutionary analyses across this group.

## Methods

### Sample collection

For this work, specimens from three *Cobitis* species (*Cobitis elongatoides*, *C. taenia*, and *C. tanaitica*) were required. All of these individuals, as well as their investigated hybrids, belonged to laboratory strains originated from natural populations, that were contained at breeding facilities at the Institute of Animal Physiology and Genetics of the Czech Academy of Sciences (see Supplementary Table S1 for detailed individual information and Figure 1 for collection distribution). The specimens were categorized into taxonomic units using published microsatellite markers and their ploidy determined by flow cytometry and karyotype verified by standard cytogenetic means as described in (Janko et al., 2012).

### RNA and DNA isolation and sequencing

A phenol/chloroform extraction protocol (Sambrook & Russell, 2006) was used to extract DNA from ∼1 g of skeletal muscle for Oxford Nanopore sequencing from one individual per species and Illumina sequencing from a second individual. DNA quality and quantity was assessed using a Qubit double-stranded DNA HS Assay Kit (Invitrogen, Thermo Fisher Scientific), agarose gel electrophoresis and an Agilent Bioanalyzer 2100 (Agilent Technologies).

For HiC, a spleen from the same specimen used for ONT was dissected and sent on dry ice to the Dovetail (*C. taenia*) and IAB (*C. elongatoides, C. tanaitica*) companies for the OMNI-C library construction.

In addition, we isolated gDNA from 122 male and female specimens of *C. taenia* and *C. elongatoides* to use for sex specific marker analysis with DNAeasy Blood&Tissue kit (Qiagen). The isolates were pooled equimolarly into four pools per species reflecting their sex and geographical origin (see Supplementary Table S1) and sequenced with Illumina 150bp paired end sequencing in IAB company.

To obtain mRNA data to be used in combination with published data (Bartoš et al., 2019) for gene annotation, mRNA was extracted from brain and gonadal tissues from several individuals of *C. elongatoides* and *C. taenia* (two easily accessible species) using the TRIzol protocol (Rio et al., 2010). Libraries were then prepared using the Lexogen SENSE Total RNA-Seq Library Prep Kit and sequenced on a NextSeq 550 with a read length of 75 bp in paired-end mode.

### Genome assembly

The Oxford Nanopore sequencing that forms the basis of the three genome assemblies were based on a single male individual from each species. The first *C. taenia* assembly was initially performed using short reads and *de Brujn* assemblers. To minimize misassemblies, we assembled the short reads with ABySS v2.0 (Jackman et al., 2017) and SOAPdenovo v2.04 (D. Li et al., 2015) and split the contigs at positions where the two assemblies disagreed. The contigs from the initial consensus assembly were combined with the nanopore reads and assembled using Flye4 v2.9.1 (Kolmogorov et al., 2019), followed by Nanopolish v0.13.1 (Loman et al., 2015). The assembly was then improved by two runs of Pilon v1.24 (Walker et al., 2014) using the Illumina sequencing data. Because we had reads from several individuals, we normalized the number of reads per individual and mapped them on the polished assembly using bwa v0.7.17 (H. Li, 2013) before variant calling with Bcftools v1.10.2 (Danecek et al., 2021). Each SNP was then modified to the major allele if necessary (custom script). This primary assembly (N50 ∼150 Kbp) was then scaffolded to a chromosomal level using Hi-C data while following the Juicer-3D-DNA pipeline v201008 (Dudchenko et al., 2017). For the genome assemblies of *C. tanaitica* and *C. elongatoides*, only flye4 with nanopore reads was used to create the primary assemblies.

### Chromosome-level De novo assembly of Cobitis genomes

Using Juicer-3D-DNA pipeline, Hi-C reads for the three species were mapped against their respective fragmented draft genome using Burrows-Wheeler Aligner (bwa) (H. Li, 2013). Reads with a MAPQ < 30 were automatically discarded. Based on the contact frequencies, 3D-DNA was run with default parameters to construct the final superscaffolds (Dudchenko et al., 2017). Final assembly stats were calculated with the script stats.sh included in the sequence-analysis package named BBmap (Bushnell, 2014). Hi-C matrices were built at 500 Kbp resolution by remapping Hi-C reads against the final assembly using Juicer with default parameters. Small scaffolds (< 30 Mbp) were discarded and only chromosome-level superscaffolds, organized by size, were included. Lastly, matrices were normalized, corrected, and plotted using ‘hicNomalize’, ‘hicCorrect’ and ‘hicPlotMatrix’ from HiCExplorer (v.3.7) (Wolff et al., 2020). First eigenvector values were calculated using the tool ‘fanc compartments’ from the HiC analysis tools package, FAN-C (v0.9.1) (Kruse et al., 2020). Moreover, topologically associated domains (TADs) were detected with the tool ‘hicFindTADs’ from HiCExplorer (v.3.7) (Wolff et al., 2020). For both analyses, normalized 50 Kbp matrices were employed as input.

### Gene annotation

*Ab initio* gene prediction was performed by Dovetail as follows: repeat families found in the genome assembly of *Cobitis taenia* were identified *de novo* and classified using the software package RepeatModeler v2.0.1 (Flynn et al., 2020). RepeatModeler depends on the programs Recon v1.08 (Bao & Eddy, 2002) and RepeatScout v1.0.6 (Price et al., 2005) for the *de novo* identification of repeats within the genome. The custom repeat library obtained from RepeatModeler was used to discover, identify and mask the repeats in the assembly file using RepeatMasker v4.1.0 (Smit et al., 2013). Coding sequences from *Triplophysa tibetana*, *Astyanax mexicanus,* and *Danio rerio* were used to train the initial *ab initio* model for *C. taenia* using Augustus v2.5.5 (Stanke, 2012). Six rounds of prediction optimisation were done with the software package provided by Augustus. The same coding sequences were also used to train a separate *ab initio* model for *Cobitis taenia* using Snap v2006-07-28 (Korf, 2004). RNAseq reads (Supplementary Table S1) were mapped onto the genome using the STAR aligner software v2.7 (Dobin et al., 2013) and intron hints generated with the bam2hints tools within the Augustus software (Keller et al., 2011; Stanke et al., 2006, 2008). Maker (Campbell et al., 2014), Snap and Augustus (with intron-exon boundary hints provided from RNA-Seq) were then used to predict genes in the repeat-masked reference genome. To help guide the prediction process, Swiss-Prot peptide sequences from the UniProt database (The UniProt Consortium et al., 2025) were downloaded and used in conjunction with the protein sequences from *T. tibetana*, *A. mexicanus,* and *D. rerio* to generate peptide evidence in the Maker pipeline. Only genes that were predicted by both Snap and Augustus were retained in the final gene annotation. To help assess the quality of the gene prediction, AED scores were generated for each of the predicted genes as part of the Maker pipeline. Genes were further characterised for their putative function by performing a Blast (Altschul et al., 1990) search of the peptide sequences against the UniProt database. tRNA were predicted using the software tRNAscan-SE v2.05 (Chan et al., 2021).

Initial gene prediction metrics were 35,082 genes. A completeness analysis showed that only 74.9% of BUSCO genes were complete, 5.9% were partially present and 19.2% were missing. This was lower than the 95% of BUSCO genes found by BUSCO if the analysis was done directly on the genome.

The annotation was improved by extending the transcripts and generating new ones using StringTie v2.2.1 (Shumate et al., 2022), followed by transdecoder v5.7.1 (Haas, 2024). We used to publish as well as newly obtained mRNA data from muscle, liver, gonad, and spleen tissue from *C. taenia* and *C. elongatoides* for annotation (Supplementary Table S1) (Bartoš et al., 2019; Janko et al., 2018). Mapping was done by STAR v2.7.10b. Both Dovetail annotation and stringtie annotation were merged using AGAT (Jacques Dainat et al., 2024). The BUSCO genes were improved to 94.6% completeness, with 1.8% fragmented and 3.6% missing genes.

Because our repetitive element annotation (next paragraph) differs from the masking used for *ab initio* gene prediction by Dovetail, we deleted genes that were identified as being repetitive elements. The final gene set from *C. taenia* was copied to *C. tanaitica* and *C. elongatoides* by Lliftoff (Shumate et al., 2022).

### Repetitive element annotation

In order to identify and annotate the repetitive elements, initial consensus sequences were generated using the Dfam TETools container v1.87 (Storer et al., 2021) (running on Docker 24.0.5), which packages RepeatModeler v2.0.5 and RepeatMasker v4.1.5 together with Dfam 3.7 (curated portion only).

We ran three RepeatModeler runs on each of the base genome assemblies (including the unplaced contigs). The resulting consensus sequences from the three species were then combined with curated families from Dfam v3.7 to form a single library.

To remove redundancy in the resulting library, we used Blastn v2.11.0+ and compared the library against itself with a word size of 20 and a minimum percentage identity of 95%. Overlapping sequences were either joined to form a new consensus or one of them shortened to remove the overlap. This was run iteratively until there were no more segments to remove. RepeatMasker was run on each genome with this reduced repeat library, which was further refined by removing portions of each sequence in the library which only aligned to the genomes once. Finally, RepeatMasker was run on each assembly using the library from the secondary refinement.

### Identification of structural variants

The assembled genomes of *C. elongatoides* and *C. tanaitica* were mapped against the *C. taenia* reference genome using minimap2 v2.24 (Li, 2018) (parameters -ax asm10 --eqx). Homology between species was checked using dot plots generated by DGennies (Cabanettes & Klopp, 2018). SyRI (Goel et al., 2019) was used to distinguish syntenic and rearranged blocks and to identify structural variants. SyRI was run independently for *C. elongatoides* and *C. tanaitica*, both analyses used the *C. taenia* genome assembly as their reference. In order to have the same number of chromosomes for all species (a requirement by SyRI identification software), Ch01A and Ch01B of *C. elongatoides* and *C. tanaitica* were combined using a 1 Kbp long spacer prior to the analyses and the coordinates of the identified structures were then transferred back to Ch01A and Ch01B.

Due to SyRI producing many short tandem events for translocations and duplications rather than one long rearrangement, all neighbouring blocks of length over 5 Kbp of the same structure type and orientation were merged to be considered a single rearrangement. This results in a more parsimonious set of changes in chromosomal structure.

Syntenic blocks, inversions, duplications, inverted duplications, translocations and inverted translocations were visualized using NGenomeSyn (He et al., 2023). For visualization purposes, only structures spanning more than 5 Kbp were considered. Gene synteny analysis was produced using python JCVI package v1.4.16 (Tang et al., 2024). Intersections of repeat annotations and indels were produced using Bedtools v2.31 (Quinlan, 2014).

### Sex Chromosome identification and validation through candidate loci PCR

Four pooled DNA samples for *C. elongatoides* and *C. taenia*, along with six individual *C. tanaitica* samples were aligned to their respective reference genomes using bwa v0.7.17 (H. Li, 2013). Pooled DNA samples were split according to sex and geographic origin of the samples, i.e. two rough geographic groupings split into two sexes resulting in the 4 pools for *C. elongatoides* (altogether 25 males and 45 females) and *C. taenia* (altogether 20 males and 32 females) respectively (see Figure 1 and Supplementary Table S1). SNPs were called from the aligned data using Gatk v4.2.3.0 (McKenna et al., 2010). Samtools v1.19.2 (Danecek et al., 2021) and Bedtools v2.31.1 (Quinlan & Hall, 2010) were used to calculate the coverage across each genome assembly and the concentration of sex-specific SNPs.

Y-chromosome specific regions were identified in the *C. elongatoides* and *C. taenia* genomes, where pooled female reads had zero coverage and pooled male reads had at least 30% of their average genomic coverage. X-chromosome specific regions were identified in *C. elongatoides* where the pooled female reads had twice the coverage as pooled male reads and within 20% of the average genomic coverage. Primers were designed to these regions, which included sections of Ch01A in *C. elongatoides* and Ch05 in *C. taenia*. In the case of *C. elongatoides*, where the identified Y-chromosome regions were larger and more numerous, PCR primers were designed with their entire range within the male specific regions. In *C. taenia*, due to a smaller relevant region, only one primer from each pair was designed within the male specific region, while the other primer was placed on the flanking regions. To evaluate the efficacy and specificity of our primers, we conducted PCR reactions (see Supplementary Table S2 for conditions) with DNA from males and females of both species (see Supplementary Table S1 for details on the individuals used). Gel electrophoresis (0.8% agarose; 80 V; TAE buffer) was performed after the PCR to confirm the amplification of single products of the expected size and to verify that the bands only appeared in expected individuals. Detailed information of three confirmed primers can be found in Table 1.

**Table 1.**
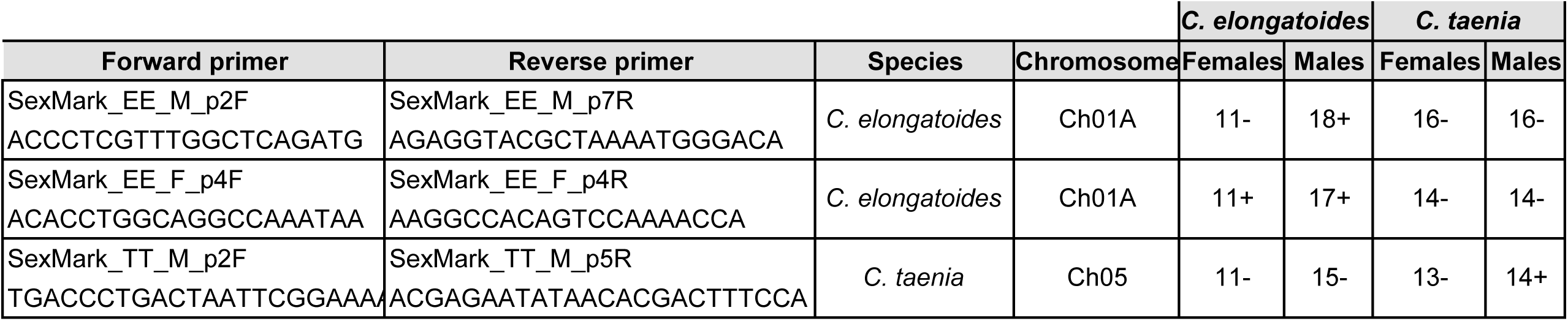
Primers used for PCR amplification and the number of tested individuals of *C. elongatoides* and *C. taenia*. A minus sign (-) indicates no detectable signal in electrophoresis after PCR amplification, while a plus sign (+) indicates the presence of a strong band.

### Candidate Sex determining genes

To identify potential candidate sex-determining genes in the three *Cobitis* species, we compiled a list of candidate actinopterygian master sex determining genes based on the literature (see Supplementary Table S3). The presence and genomic location of these genes were then determined using the gene annotation of each species and verified with Tblastn (see Supplementary Table S4).

### Cytogenetics

#### Mitotic and meiotic chromosome preparation

In order to visualize chromosomal pairing, adult *C. elongatoides*, *C. tanaitica,* and *C. taenia* males and females as well as *C. elongatoides* x *C. taenia* hybrid males were injected with 0.1% colchicine solution (1 ml/100 g of body weight). Mitotic and meiotic metaphase chromosome spreads were obtained from kidneys and testes according to previously published protocols (Majtánová et al., 2016; Marta et al., 2020). Briefly, kidneys and testes were removed, dissected in 0.075 M KCl to release cells and treated hypotonically for 30 min at room temperature. After centrifugation, cells were fixed in freshly prepared fixative methanol: acetic acid (3:1) and washed twice in a new portion of fixative. The fixed cell suspension was then dropped onto slides. Mitotic and meiotic metaphase chromosomes were initially stained with Giemsa to assess chromosomal number and morphology.

### Single chromosome Oligo-FISH probe design and Chromosomal painting

Oligomers specific to Ch01A, Ch01B, Ch05, and Ch20 were designed against the *C. taenia* assembly using Chorus software (T. Zhang, 2024). Concerning different chromosome lengths, one set of 27,000 oligomers (45-mers) was designed to visualize Ch20, and two sets of oligos were designed to detect the longer Ch01a, Ch01b, and Ch05 (Table 2). The chromosome regions with very low oligo densities were omitted in the final probe datasets (Supplementary Table S5) that are available under request. The final probe sets were synthesized as myTAGs® Labeled Libraries (Daicel Arbor Bioscience) and directly used for FISH experiments.

**Table 2:** Final genome assembly statistics for the three *Cobitis* species.

| Species | *C. taenia* | *C. tanaitica* | *C. elongatoides* |
| --- | --- | --- | --- |
| Number of Contigs | 7.2k | 25.8k | 28.9k |
| Cumulative Length (Gbp) | 1.63 | 1.71 | 1.82 |
| N50 (Mbp) | 53.2 | 51.3 | 43.0 |
| L50 | 13 | 14 | 17 |
| Number of Chromosomes | 24 | 25 | 25 |
| assembled into | 85.8% | 69.8% | 68.4 |

Commercially synthesized oligoprobes to chromosomes 5 (Ch05) and 20 (Ch20) as well as for the long (Ch01A) and short (Ch01B) arms of *C. taenia* chromosome 1 (Arbor Biosciences) were used for hybridization procedure. Oligoprobes to Ch20 and Ch01A were labelled with biotin, and oligoprobes to Ch05 and Ch01B were labelled with digoxigenin. Prior to hybridization, chromosome slides were incubated with 0.01% pepsin/0.01 M HCl at room temperature for 10 min and fixed with 2% paraformaldehyde (PFA) for 10 min. For two-color FISH we mixed oligoprobes to Ch05 and Ch20 or Ch01A and Ch01B (50 ng of each probe per slide) with 20 ul of hybridization mixture (50% formamide, 10% dextran sulfate, 2× ЅЅС, and 500 ng of salmon sperm DNA (Sigma-Aldrich). Probes were denatured at 86 °C in the heating block for 10 min and then put on ice. Slides with mitotic or meiotic chromosomes were denatured in 75% formamide at 74 °C for 5 min, dehydrated in an ice-cold series of ethanol (70%, 80%, 96%) and dried prior to denatured probe application. After hybridization overnight at room temperature, slides were washed three times with 0.2x SSC for 5 min at 42°C and 2× SSC for 5 min at room temperature. The biotin and digoxigenin labelled probes were detected using streptavidin-AlexaFluor 488 (Invitrogen) and anti-digoxigenin-rhodamine (Invitrogen), respectively. After washings in 4× SSC with 0.1% Tween at 44 °C for 5 min with shaking, slides were dehydrated in ethanol series (70%, 80% 96%), air dried, and mounted in Vectashield medium containing DAPI (1.5 mg/ml) (Vector).

### Wide-field and fluorescence microscopy

Mitotic and meiotic chromosomes with chromosomal painting were inspected using Carl Zeiss Axio Imager.Z2 and Provis AX70 Olympus microscopes equipped with standard fluorescence filter sets. Microphotographs of chromosomes were captured by CCD camera (DP30W Olympus) using Olympus Acquisition Software and CoolCube 1 using Metasystem platform for automatic search, capture and image processing. Microphotographs were finally adjusted and arranged in Adobe Photoshop, CS6 software.

## Results

### Genome assembly

The assembly process of *C. taenia* involved multiple sequencing technologies and scaffolding techniques to achieve the final high-quality genome assembly from a combination of deep coverage with short reads and lower coverage with long ONT reads. The still fragmented initial assembly (N50 ∼150 Kbp) was subsequently used to generate superscaffolds with Hi-C contact reads (Figure 2A, Table 2, see the Supplementary Table S6 for comparison of assemblies before and after Hi-C interaction mapping).

**Figure 2.**
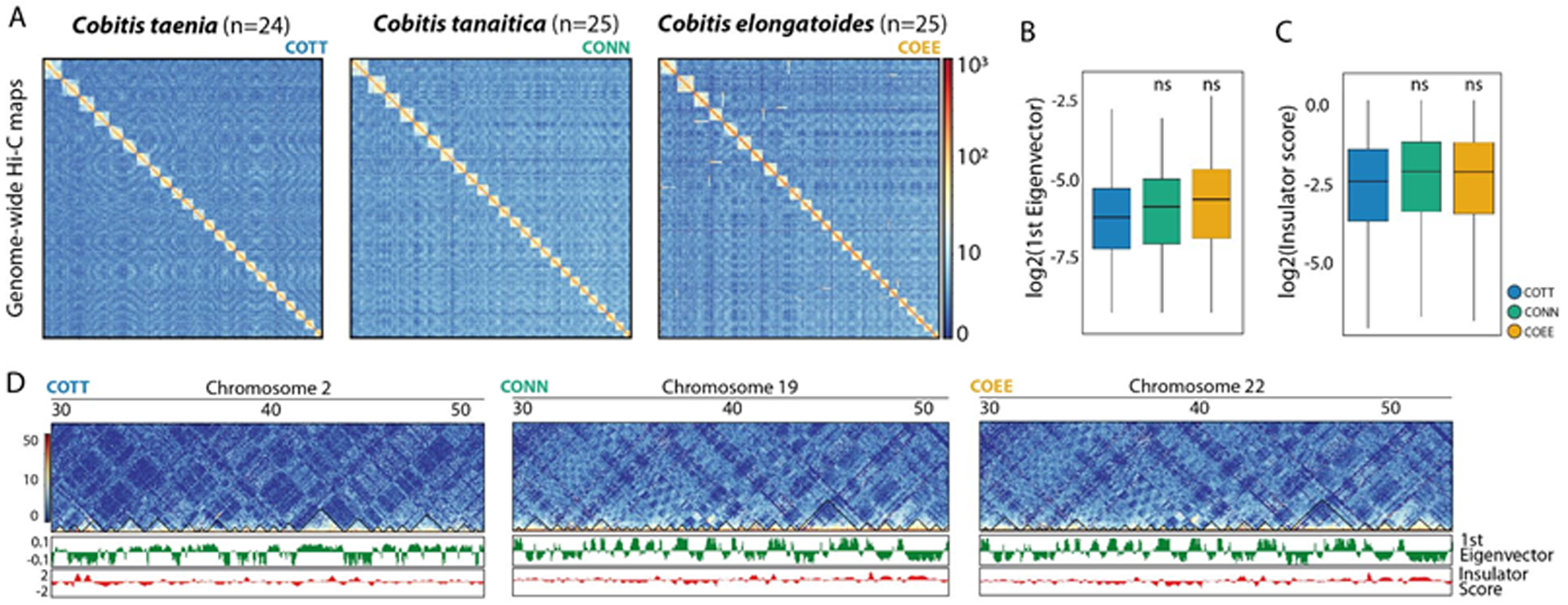
*Cobitis* genomes higher-order chromatin organization. (A) Genome-wide Hi-C contact maps. Contact maps represent 500 Kbp resolution Hi-C matrices obtained using the final assembly as a reference. For the three species clear interacting blocks corresponding to the expected number of chromosomes can be observed. (B) Boxplot depicting the 1^st^ Eigenvector distribution of the three species. Eigenvector +/- values are used as a proxy to determine open (A compartments) and close (B compartments) chromatin regions. The similarities in the distribution between the three species indicate similar 3D organization (two-sided t test, ns p > 0.05). (C) Boxplot showing insulator score distribution on the three species. Insulator capacity is used to determine TADs strength. Like eigenvector distribution, similarities on the insulator score reflect the same patterns of chromatin folding in the three species (two-sided t test, ns p > 0.05). (D) Region-specific 500 Kbp heatmaps, 1^st^ eigenvector and insulator score tracks in the three species. Similar tendencies can be clearly observed.

To generate high-quality genome assemblies (N50 > 40 Mbp), Hi-C contact reads were used to scaffold the initial fragmented genomes (N50 ∼150 Kbp) (Supplementary Table S6) into chromosome-level superscaffolds (Supplementary Table S7). A total of 416 (*C. taenia),* 801 (*C. tanaitica)* and 988 million read pairs *(C. elongatoides)* of Hi-C reads were employed to assemble each genome using the Juicer-3D-DNA pipeline (see methods). After filtering, a total of 109 million (*C. taenia)*, 427 million (*C. tanaitica)* and 433 million *(C. elongatoides)* unique contacts were used to assemble each species genome (Supplementary Table S8).

Ultimately, we successfully generated chromosome-level assemblies for males of the three *Cobitis* species: *C. taenia* (n=24, where “n” is the number of chromosomes in a haploid set), *C. tanaitica* (n=25), and *C. elongatoides* (n=25); Table 2, Figure 2A) with N50 values greater than 40 Mbp, indicating high-quality and well-scaffolded genomes. The final genome sizes were 1.6 Gbp for *C. taenia*, 1.7 Gbp for *C. tanaitica*, and 1.8 Gbp for *C. elongatoides*. The majority of each genome, namely 85.8% in *C. taenia*, 69.8% in *C. tanaitica* and 68.4% in *C. elongatoides*, was organized into chromosome-level superscaffolds whose numbers correspond with the described diploid numbers (2n=48 in *C. taenia* and 2n=50 in *C. elongatoides* and *C. tanaitica*) (Janko et al., 2007).

Chromosome-level scaffolds were named based on their length in the *C. taenia* genome from Ch01 (the largest) to Ch24 (the smallest) (Supplementary Table S9). This suggested chromosome nomenclature is based on chromosome scaffold size and does not correspond to previously published classifications based on chromosome morphology (e.g., Janko et al., 2007; Marta et al., 2020). In the other two species, we followed the same names and orientation for inferred homologous chromosomes. In the case of Ch01, which represents a recently fused chromosome in *C. taenia*, we labelled the two homologous chromosomes in *C. elongatoides* and *C. tanaitica*, which represent the ancestral syntenies, as Ch01A and Ch01B.

### A/B compartments and topologically associated domains (TADs)

Comparison of Hi-C matrices revealed different patterns of chromosomal interactions (Figure 2). The detection of 3D structures, such as compartments and TADs revealed similar patterns of chromosome folding in the three species (Figure 2A), mirroring previous observations in vertebrate species (Álvarez-González et al., 2022; Pérez-Rico et al., 2020). The genome-wide distribution of A/B compartments was similar across taxa. Consistently, ≈50% of the genome was detected as A compartments (“open” chromatin) and no major differences were observed on compartment strength between species (Figure 2B). Topologically associated domains showed the same trends. In the three species, domains of ≈0.8 Mbp (Supplementary Table S9) were defined with equal insulation capacity (Figure 2C, D). Overall, our results suggest a high level of structural conservation among *Cobitis* species.

### Inference of homology and Structural variants

Assembled genomes were aligned to each other in order to identify homologous chromosomes between the species. There was a clear one to one alignment pattern observable along the entire chromosomal length for 23 out of the 24 (in *C. taenia*) or 25 (in *C. tanaitica* and *C. elongatoides*) chromosomes (Figure 3A, 3B, Supplementary Figure 1). The mapping of the remaining chromosome (Chr01) of *C. taenia* was split between the two remaining chromosomes (Chr01A and Chr01B) of *C. tanaitica* and *C. elongatoides* indicating a chromosomal fusion in *C. taenia*.

**Figure 3.**
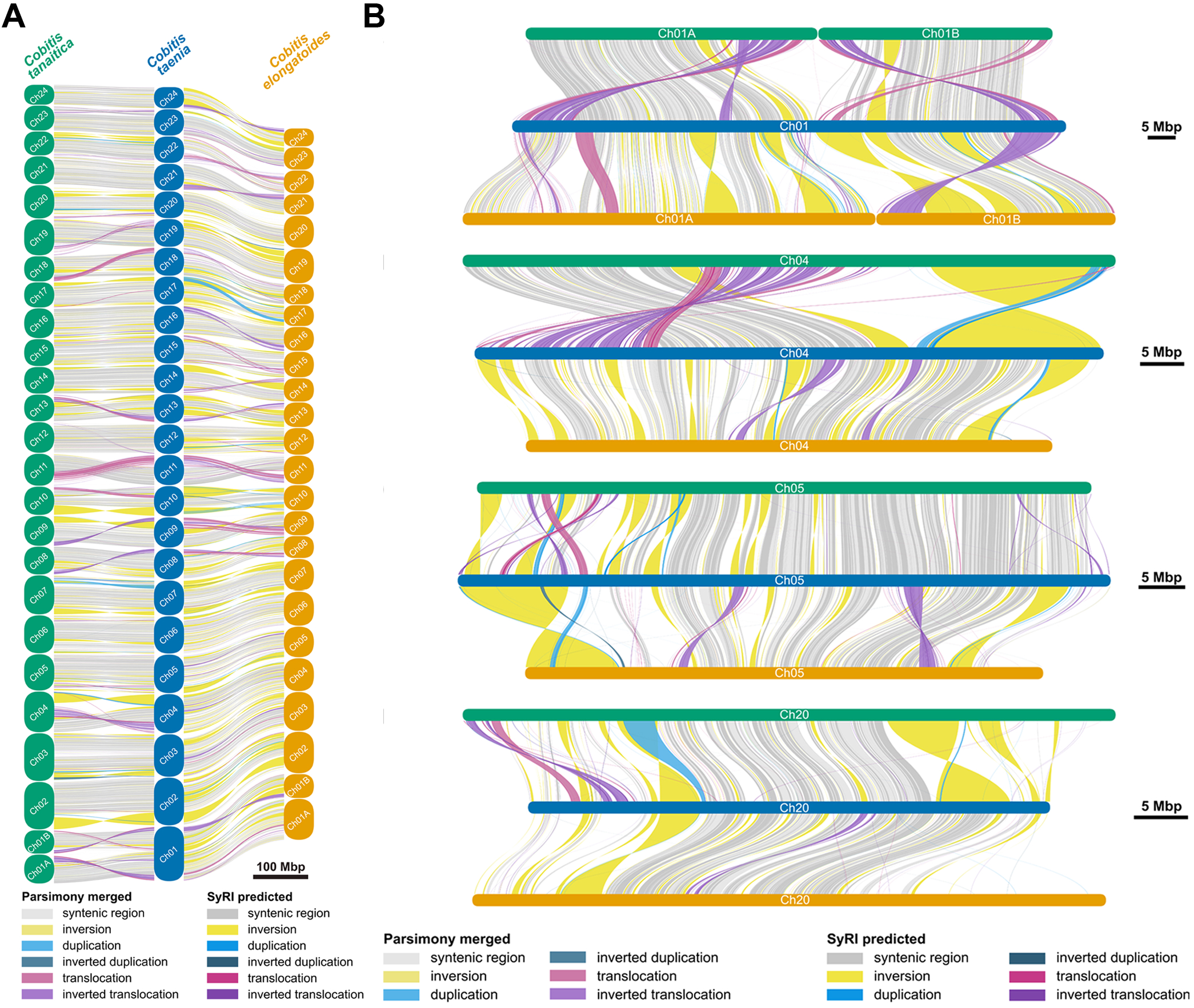
Synteny plots of homologous sequences and intrachromosomal rearrangements in three *Cobitis* genomes. A) Syntenic regions and rearranged blocks are shown along the three genomes (from left to right *Cobitis tanaitica*, *C. taenia* and *C. elongatoides*). B) Four selected chromosomes in larger detail (chromosome 1, chromosome 4, chromosome 5, chromosome 20) of the three species (from top to bottom *C. tanaitica*, *C. taenia* and *C. elongatoides*). Darker colour shades highlight the SyRI detected events while lighter shades show the results of parsimonious merging. Only blocks longer than 5 Kbp are shown.

Syntenic regions and rearrangements in the chromosomes were identified using SyRI followed by a merging procedure to ensure parsimony (see Methods). The proportion of syntenic regions between homologous chromosomes ranged from 24 to 68% in the comparison of *C. tanaitica* to *C. taenia* and from 17 to 49% in the comparison of *C. elongatoides* to *C. taenia*. We identified a total of 35,633 variants in the comparison of the *C. tanaitica* genome to the *C. taenia* genome (*C. taenia* to *C. tanaitica*) and 41,228 variants in the comparison of *C. elongatoides* to the *C. taenia* (*C. taenia* - *C. elongatoides*) (Supplementary Table S11). Among the longest rearrangements, we detected 53 and 66 inversions spanning over 1 Mbp in the *C. taenia* - *C. tanaitica* and *C. taenia* - *C. elongatoides* comparisons, respectively. Other types of rearrangements, such as duplications and translocations, were however scattered in an excess of adjacent shorter events when they would often be better explained by a single larger event. For the intrachromosomal rearrangements, we remedied this with our parsimony merging procedure (see Methods) and got to a total number of 69 (in the *C. taenia* - *C. tanaitica* comparison) and 116 (in the *C. taenia*

- *C. elongatoides* comparison) rearrangements spanning over 1 Mbp (Figure 3A, Supplementary Table S11). These extra-long rearrangements include large translocations in Ch11 and Ch04 in *C. tanaitica* having approximately one-third of the chromosome spanned by inverted translocations (Figure 3B).

Identified insertions and deletions spanned a total length of 7.6 Mbp and 2.8 Mbp in the *C. taenia* - *C. tanaitica* and *C. taenia* - *C. elongatoides* comparisons respectively. Of these indels, approximately 73% were composed of repeats in the *C. taenia* - *C. tanaitica* comparison and 71% in the *C. taenia* - *C. elongatoides* comparison. Among the most represented classes of repeats among the indels were DNA hAT-Ac elements (9% in *C. taenia*

- *C. tanaitica*, 4.8% in *C. taenia* - *C. elongatoides*) and LTR Gypsy elements (10.6% in *C. taenia* - *C. tanaitica*, 12.4% in *C. taenia* - *C. elongatoides*).

The intrachromosomal rearrangements we detected were confirmed using gene synteny. We also observed several interchromosomal events, 12 in the *C. taenia* - *C. tanaitica* comparison, 50 in the *C. taenia* - *C. elongatoides* comparison and 100 in the *C. tanaitica* - *C. elongatoides comparison* (Supplementary Figure 2). These interchromosomal events span on average 8 genes in *C. taenia - C. tanaitica*, 11 genes in *C. taenia* - *C. elongatoides* and approximately 9 genes in *C. tanaitica - C. elongatoides*. The longest detected rearrangements (spanning over 40 genes) from the *C. taenia* - *C. elongatoides* comparison are located on Ch05, Ch19, Ch22, and Ch23 in *C. elongatoides* and Ch02, Ch03, Ch10, and Ch22 in *C. taenia*. Interchromosomal rearrangements between these pairs of chromosomes are also the longest ones in the *C. tanaitica* - *C. elongatoides* comparison.

### Gene annotation

For *C. taenia* we identified 31,513 genes, generating 95,721 different mRNAs. BUSCO analysis using actinopterygii_odb10 (3 640 BUSCOs) shows 94.6% coverage (BUSCO v5.8.2, ODB v10, hmmsearch v3.4). Both other genomes give slightly worse results, with *C. elongatoides* having 30,420 genes (89,355 transcripts) with a BUSCO coverage of 81.5% and *C. tanaitica* having 30,341 genes, 90,600 transcripts and a BUSCO coverage of 85.4%. The number of genes per chromosome and the proportion of scaffolded versus unscaffolded parts of the genome are given in Supplementary Table S9.

### Repetitive element annotation

Overall, approximately 54-55% of the assembled *Cobitis* genomes were identified as being repetitive elements. This includes 9,928 repeat classes and 90 repeat families in *C. taenia*, 9,882 repeat classes and 90 repeat families in *C. elongatoides*, and 9,924 repeat classes and 91 repeat families in *C. tanaitica*, excluding simple, low complexity, and satellite repeats.

Most families are equally distributed (Figure 4A), which suggests they expanded before speciation, and they are no longer active. But several families are expanded only in one or two species, suggesting that they were active after speciation. These families could therefore be used in the future for further detection of deregulation in hybrids.

**Figure 4.**
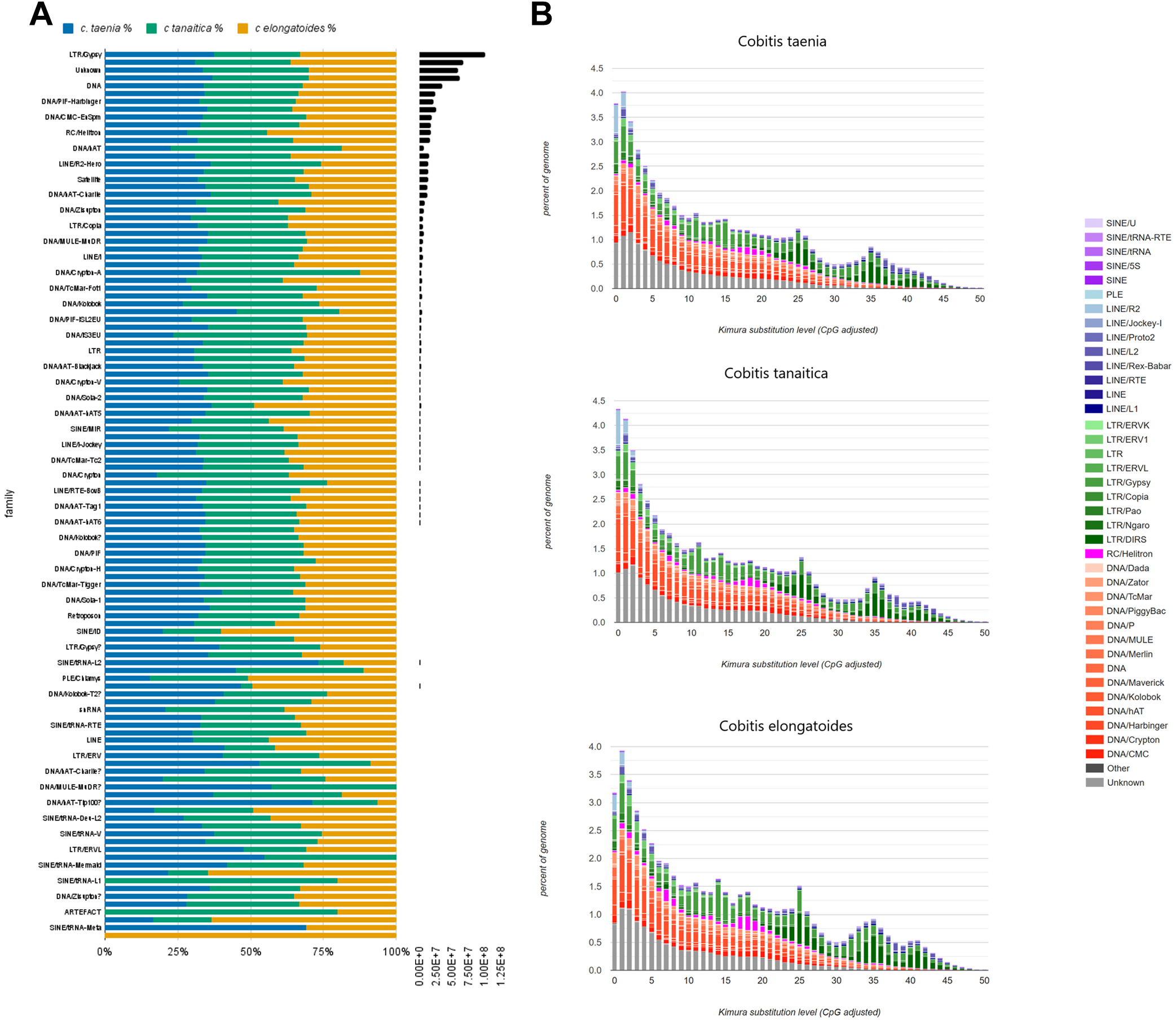
Repeatome in *Cobitis*. A) A comparison of relative distribution of individual TE families for each genome (colored by species). Black bars show the absolute lengths of each family in *C. taenia*. B) The repeat landscape plot illustrating TE accumulation history for the three *Cobitis* genomes (*C. taenia, C. tanaitica, C. elongatoides*), based on Kimura distance-based copy divergence analyses. The sequence divergence (CpG adjusted Kimura substitution level) is shown on the x-axis while the percentage of the genome represented by each TE type is on the y-axis. Transposon type is indicated by the key on the right.

A Kimura plot analysis demonstrated that the closely related *C. taenia* and *C. tanaitica* have very similar TE content with clear evidence of an on-going expansion in some families, while *C. elongatoides* differs from both with indications of a recent decline in TE activity (Figure 4B).

### Identification of sex chromosomes

Pooled genomic sequencing from 24 males and 45 females from *C. elongatoides* and 20 males and 32 females from *C. taenia* were used to calculate the sex specific coverage and SNPs across each respective reference genome. Due to the difficulty of getting *C. tanaitica* individuals, three males and three females were sequenced individually and used for the analysis. In *C. elongatoides*, this resulted in a clear signal across the whole Ch01A scaffold, which represents a separate chromosome in *C. tanaitica* and other cobitoids, but which has fused with Ch01B in *C. taenia*. Portions of this chromosome-level scaffold in *C. elongatoides* had a coverage in females equal to the average genomic coverage and half the coverage in males, consistent with an X chromosome, while other regions of the chromosome showed half the average genomic coverage in males and low coverage in females, consistent with a Y chromosome (Figure 5C). This intermixed pattern suggests that *C. elongatoides* has an X/Y system and that the assembled Ch01A scaffold represents a chimeric combination of these chromosomes’ sequence. No visible coverage differences were observed in any chromosome in *C. taenia* and *C. tanaitica* (Figure 5 A-B), suggesting that these species have, compared to *C. elongatoides*, young undifferentiated sex chromosomes.

**Figure 5.**
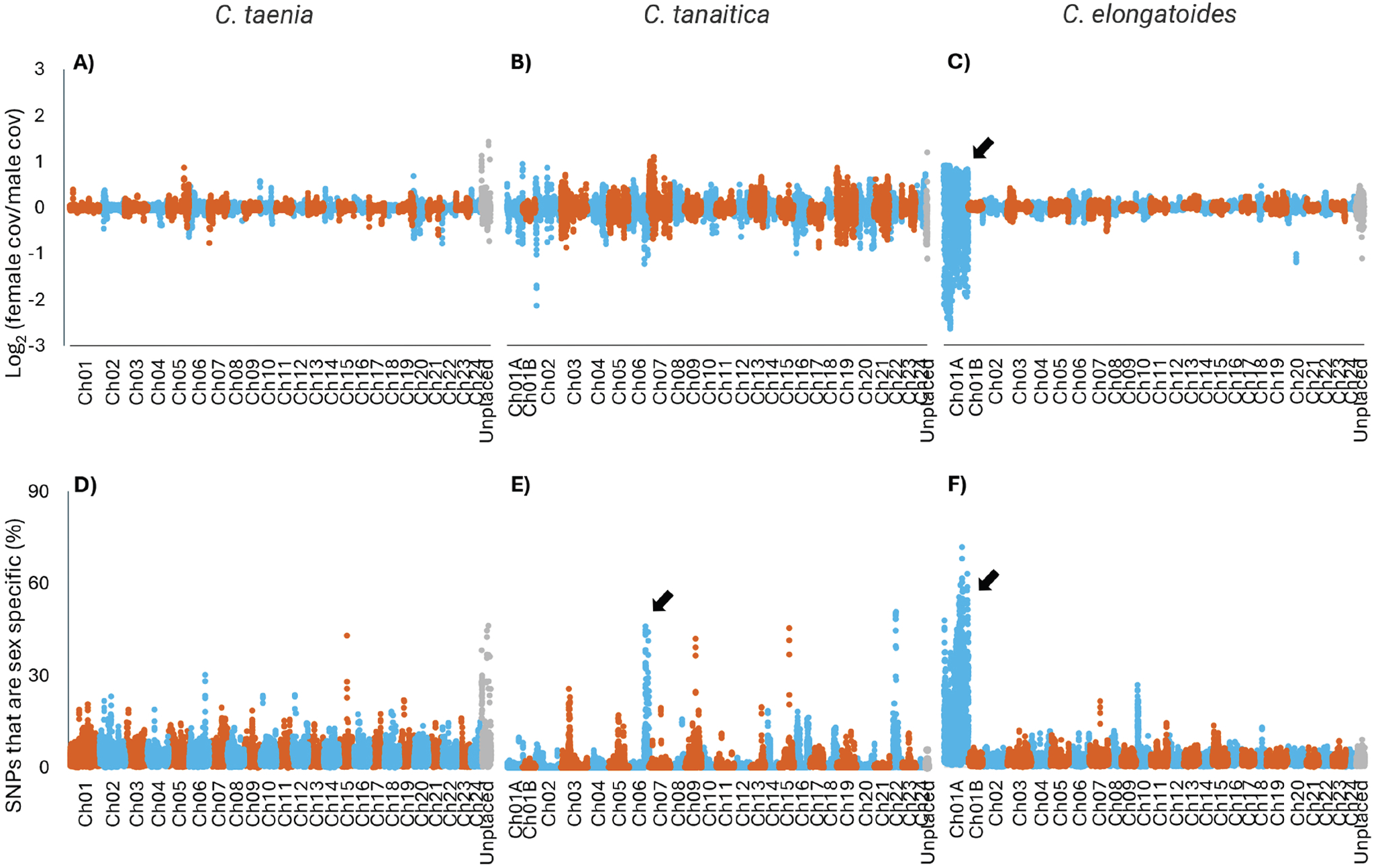
Signals of sex-linked differentiation across the three *Cobitis* genomes. Differences in coverage between male and female individuals are shown on the top row (A-C), with values representing the log2 transformed coverage in females divided by the coverage in males. The concentrations of sex-specific SNPs are shown on the bottom row (D-F) with values normalised by the number of SNPs identified in that region. Each data point represents the total across a window of 200kb of the genome, with consecutive windows starting 50kb apart. *C. taenia* (A, D) and *C. elongatoides* (C, F) are created using pooled DNA from males and females while *C. tanaitica* (B, E) is created from 3 male and 3 female individuals. Arrows point to the most promising (if any) differentiated sex regions.

Sex-specific SNPs were identified in all three species using the pooled Illumina datasets. The proportion of these sex-specific SNPs was used to locate regions of differentiation between male and female individuals. This revealed several potential sex-specific regions in *C. tanaitica*, most notably a 900kb region on Chromosome 6 (Figure 5E) as well as once again highlighting the whole of Chromosome 1A in *C. elongatoides* (Figure 5F). No concentration of sex-specific SNPs was observed in *C. taenia* (Figure 5D).

As a final attempt to identify a sex chromosome in *C. taenia*, regions were identified with zero coverage in females and a coverage greater than 30% of the average genomic coverage in males (where the Y chromosome-specific regions are expected to have an average of 50% coverage). The largest such region, which was 246bp, was found on Ch05. PCR primers were designed within this region and successfully validated using an additional 11 sexed individuals not used in the Pool-Seq data.

Primers were also designed to amplify regions predicted to be sex chromosome-specific (either X- or Y-specific) in *C. elongatoides* to use for future sex identification of pure individuals or identification of sex chromosome presence in interspecific hybrids. The PCR primers and their amplification conditions were tested on the individuals that were not used in the creation of the PoolSeq library. As a criterion for primer selection for further screening we considered that primers were (1) amplifying a single band in a gonosome-specific manner (i.e. Y-marker in males only and X-marker in both sexes), and (2) they amplified in a species-specific manner, hence, not making any product in non-target species. Finally, three sets of primers were selected, reliably diagnosing Ch01A Y- and X-specific loci in *C. elongatoides* and Ch05 Y-specific loci in *C. taenia* (Table 1). A primer set that exclusively amplified the *C. taenia* X locus was not found.

From the literature (Supplementary Table S3), 42 genes (including paralogs) were found which are master sex determination or key regulators of sex determining pathways in actinopterygians. All of these genes were mapped to chromosome-level scaffolds except four in *C. tanaitica* and nine in *C. elongatoides*, which were found on unplaced scaffolds. Notably, among these genes, *paics* (the master sex determination genes in the blue tilapia - *Oreochromis aureus*) aligned to Ch01A, the putative sex chromosome, in *C. elongatoides* (Supplementary Table S4).

Additionally, we analyzed the list of genes located on Ch01A to check for the presence of any additional genes, which were considered as linked to sex differentiation in various fish species. We detected the presence of four such genes (*bmp2b, gata4, gpatch2,* and *gopc*) which are known to play a key role in vertebrate sex determination and thus are potential candidates for the sex determination master gene in *Cobitis elongatoides*. Interestingly, *gata4* and *gpatch2* are located in a highly differentiated region of the X chromosome while *gopc* and *bmp2b* are in a moderately diverged region. *paics* is in the least diverged part. All these genes are located on the same syntenic group (autosome 20) in *Danio rerio*.

### Cytogenetic Validation of Chromosome Structures Through Mitotic Chromosome Painting

Chromosome painting was performed using probes designed using the genome assemblies. These included probes that covered the whole of Ch20, as well as probes covering parts of Ch01A, Ch01B and Ch05. These paintings confirmed the accuracy of our genome assemblies across four selected linkage groups in all three sexual *Cobitis* species. These probes, designed using all three species-specific assemblies, consistently illuminated the targeted regions, affirming the scaffolded linkage groups. Specifically, in Ch05 the probe highlighted the long arm of a large submetacentric chromosome across all species, confirming the structural integrity and assembly accuracy of this chromosome. In Ch20 the probe marked the q-arm of small subtelocentric chromosomes in *C. taenia* and *C. elongatoides*, suggesting a conserved structure in these species. Conversely, in *C. tanaitica*, the probe signal was observed in the q-arm of a small acrocentric chromosome (Figure 6), suggesting morphological differences. Finally, the application of probes for linkage groups Ch01A and Ch01B in *C. elongatoides*, when applied in *C. taenia*, showcased distinct hybridization signals on the short and long arms of the largest metacentric chromosome with its centromeric region unstained and exhibiting only DAPI signal (Figure 6). This confirmed a fusion event unique to this species. In *C. tanaitica* the Ch01A and Ch01B probes highlighted two pairs of acrocentric chromosomes and in *C. elongatoides*, two pairs of submetacentric chromosomes, once again highlighting structural differences among the species. Interestingly, even after the visualisation of Ch01A, we did not observe any difference in morphology and hybridisation signals between male and female mitotic chromosomes in all three studied sexual species, despite the possible role of Ch01A in sex determination of *C. elongatoides* and the identified divergence between Y- and X-specific sequences. This suggests that differentiation of the XY chromosomes is limited to nucleotide substitutions and other small rearrangements rather than large structural changes.

**Figure 6.**
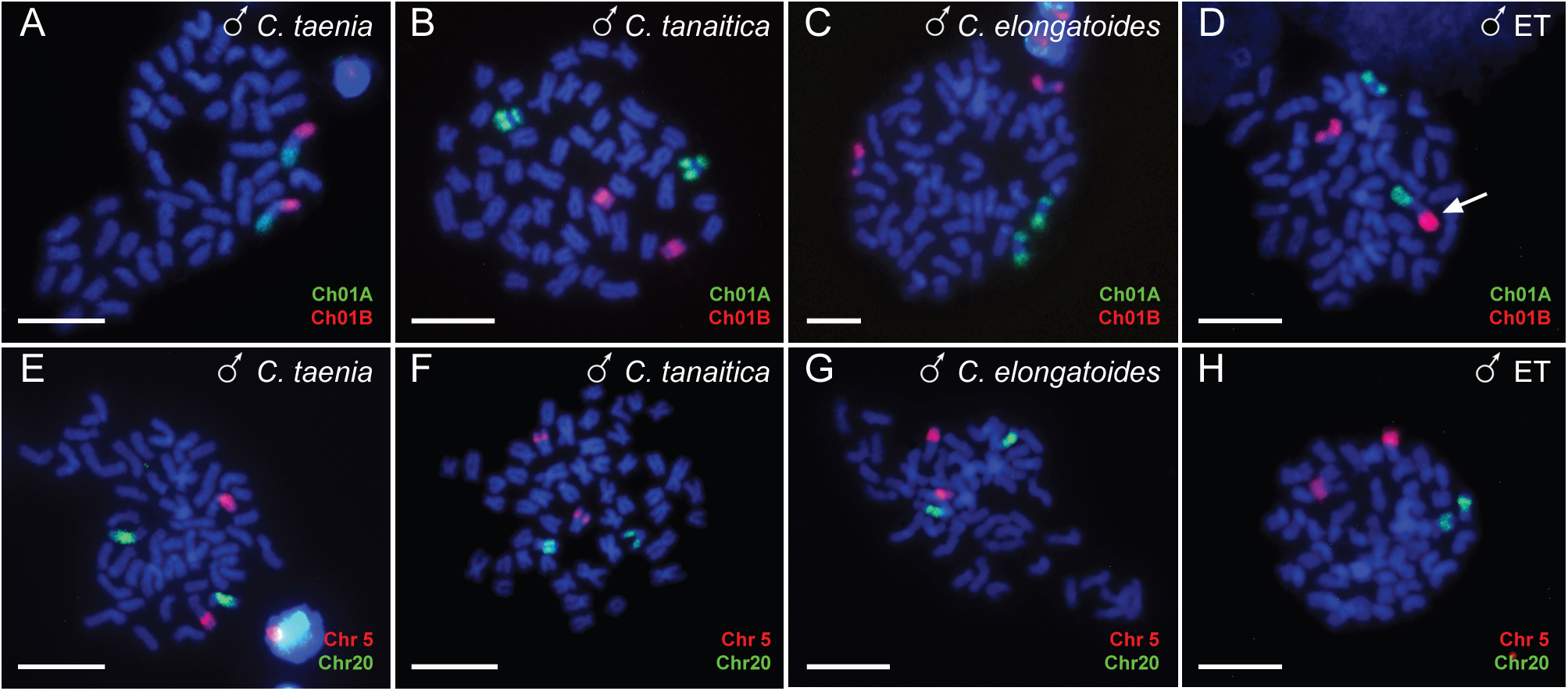
Chromosome painting of selected chromosomes. Ch01A and Ch01B (A-D) and Ch05 and Ch20 (E-H) are shown on mitotic metaphases of *C. taenia* (A, E), *C. tanaitica* (B, F), *C. elongatoides* (C, G) and diploid ET hybrid (D, H) males. Ch01A (green) and Ch01B (red) are located on different arms of the largest metacentric chromosome in *C. taenia* (A). In *C. tanaitica,* signals appeared on two pairs of acrocentric chromosomes (B) and in *C. elongatoides*, they were located on two pairs of submetacentric chromosomes (C). Small submetacentric chromosome stained by Ch01A represents the sex chromosome of *C. elongatoides* (C). In diploid ET hybrid, both signals were detected on one metacentric chromosome of *C. taenia* (pointed by arrow) and two submetacentric chromosomes of *C. elongatoides* (D). Chromosome painting of Ch05 showed signals on the long arm of a large submetacentric chromosome across all species (E-G) and in diploid hybrid (H). Chromosome painting of Ch20 (green) was detected in the q-arm of a subtelocentric chromosome in *C. taenia* (E) and *C. elongatoides* (G) and corresponding chromosomes in the diploid hybrid (H) but locates in a q-arm of a subtelocentric chromosome in *C. tanaitica* (F). Chromosomes are stained by DAPI (blue). Scale bar = 10 µl.

In diploid elongatoides-taenia (ET) hybrids, chromosome painting verified the presence of orthologous Ch05 and Ch20 corresponding to those identified in *C. elongatoides* and *C. taenia*. Further, the Ch01A and Ch01B probes revealed the fused *C. taenia*’s chromosome Ch01 alongside the distinct submetacentric chromosomes from *C. elongatoides* (Figure 6).

### Cytogenetic Exploration of Meiotic Chromosome Spreads in Hybrids and Pure Species

Chromosome painting with Ch05 and Ch20 probes applied to meiotic metaphase I spermatocyte spreads showed, in all three parental species, the presence of one larger bivalent corresponding to Ch05 homologs and a smaller bivalent corresponding to Ch20 homologs (Supplementary Figure 3). In addition, meiotic metaphase I of *C. taenia* showed both Ch01A and Ch01B hybridization signals on the single largest bivalent, while *C. elongatoides* and *C. tanaitica* exhibited two distinct small bivalents corresponding to Ch01A and Ch01B paired homologs, corroborating the mitotic chromosome data (Supplementary Figure 3).

To understand how the pairing patterns proceed in interspecific hybrids, we applied these chromosome-specific probes to 526 meiotic metaphase I spermatocytes from two diploid elongatoides-taenia (ET) hybrid males (Figure 7A-C). These males were derived from crossing of *C. taenia* mothers and *C. elongatoides* fathers and were now confirmed to have inherited the Y gametolog of Ch01A from *C. elongatoides* via positive PCR amplification signal with the Y-linked primer pair and negative amplification for the X-linked primer pair (see Table 1).

**Figure 7.**
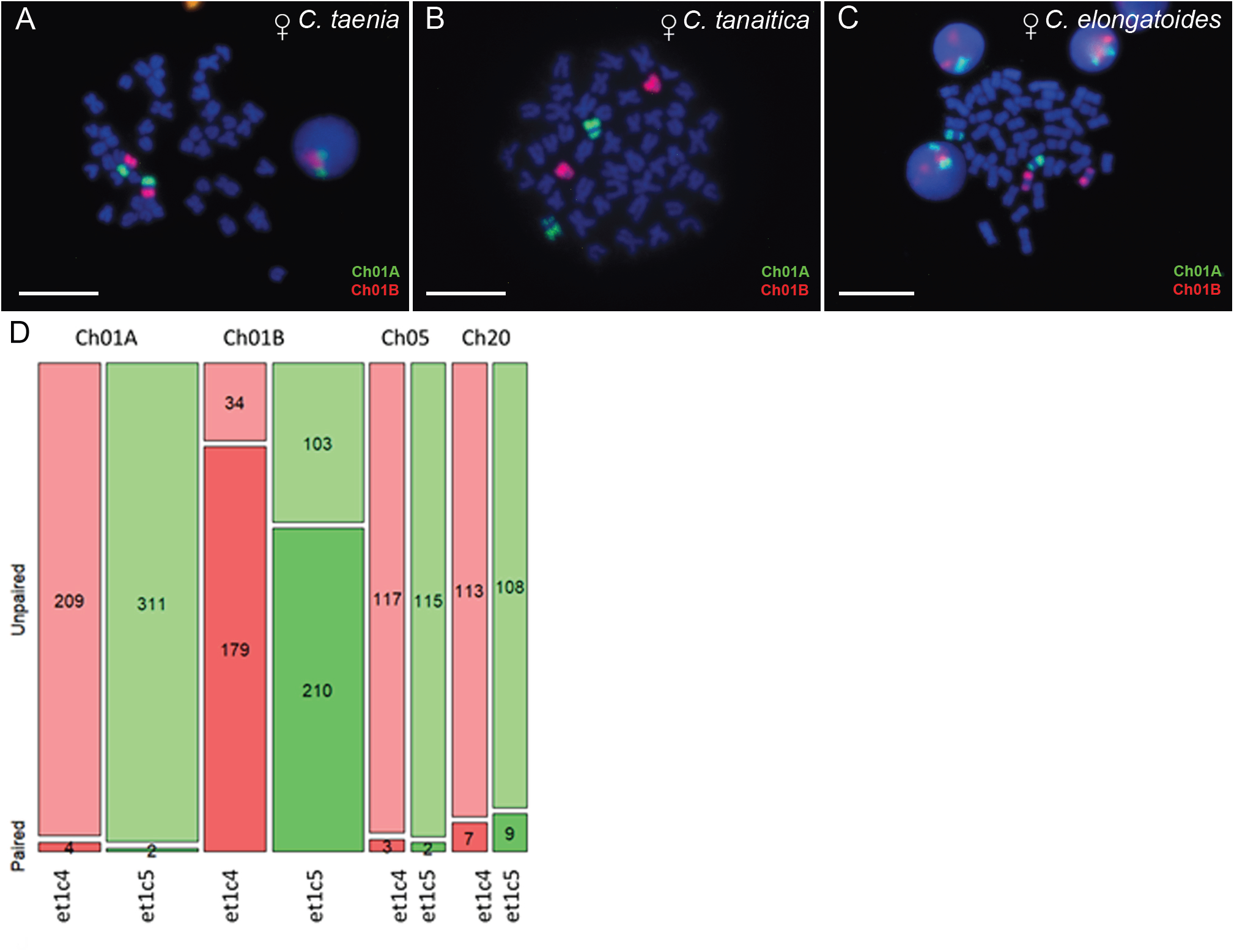
Chromosome pairing in hybrid males. (A-C) Chromosome painting of Ch05 and Ch20 (A) as well as Ch01A and Ch01B (B, C) on meiotic metaphases of diploid hybrid males. Probes for Ch05 and Ch20 were located on *C. taenia* and *C. elongatoides* chromosomes that did not show pairing between orthologs and existed as univalents (A). Probes for Ch01A and Ch01B hybridized to the *C. taenia* chromosome and two small chromosomes of *C. elongatoides* (B, C). Chr01A of *C. elongatoides* usually existed as univalent (B, C). In some spermatocytes, *C. elongatoides* chromosome hybridizing with probe for Chr01B showed pairing with homologous arm of Chr01B of *C. taenia* (B), while in other spermatocytes, there was no pairing between part of the Chr01B of *C. taenia* and chromosome Chr01B of *C. elongatoides* (C). Chromosomes are stained by DAPI (blue). Scale bar = 10 µl. D) Mosaic plot showing pairing success of four chromosomes (Ch01A, Ch01B, Ch05, Ch20) in two diploid hybrid males. Each chromosome is represented by two adjacent bars (one per individual), with bar height normalized to 100%. The lower shaded section indicates the proportion of cells where the chromosome formed a bivalent, while the upper section represents univalents. Numbers within each section show absolute counts. Bar width reflects the total number of spermatocytes analyzed per chromosome and individual. Ch01B exhibited the highest pairing frequency, whereas Ch05, Ch20, and especially Ch01A were mostly unpaired.

Despite differences in the total number of metaphases inspected for each individual and chromosome (see Figure 7D for exact counts), statistical analysis revealed significant differences in pairing success among the four investigated chromosomes. A generalized linear model (GLM) with a binomial error structure was used to test the effects of chromosome identity, individual, and their interaction on pairing success (response variable: paired vs. unpaired). The model showed that chromosome identity had a highly significant effect on pairing success (p < 0.0001), indicating consistent interchromosomal differences in pairing likelihood.

Post-hoc pairwise comparisons, adjusted for multiple testing using the Bonferroni correction, confirmed that Ch01B paired significantly more frequently than all other chromosomes (p < 0.0001 for all pairwise contrasts; Figure 7D). Ch05 and Ch20 bivalents were observed in only a small proportion of cells, with both chromosomes showing significantly lower pairing likelihoods compared to Ch01B (p < 0.0001 for both comparisons). Ch01A, the putative Y gametolog derived from *C. elongatoide*s, exhibited the lowest pairing rates overall, appearing primarily as univalents (Figure 7D).

While these interchromosomal differences were consistent across both individuals, significant interindividual differences were observed for Ch01B. A separate GLM for Ch01B revealed that its pairing success was significantly lower in one hybrid male (*et1c5*) compared to the other (*et1c4*; p = 0.0006). However, no significant interindividual differences were detected for the other chromosomes, suggesting that variability in pairing dynamics for Ch01B might reflect unique characteristics of this chromosome or its interaction with individual-specific factors.

These findings underscore the variability in chromosomal pairing success, with Ch01B showing the greatest likelihood to form bivalents, while the Y gametolog of Ch01A from *C. elongatoides* rarely pairs with its ortholog from *C. taenia*. This variability likely contributes to the observed chromosomal incompatibility and reproductive barriers in these hybrids.

## Discussion

### Filling the taxonomical gap in Chromosome Level assemblies

Recent advances in long read sequencing and chromatin capture technologies have substantially improved the feasibility of chromosome-level genome assemblies for non-model organisms. We took advantage of this trend to generate three high quality *Cobitis* genome assemblies. While assemblies created with these technologies represent a major improvement over short read assemblies, they can still struggle with genomic features such as repetitive regions, centromeres, telomeres, and structural variants. This can lead to fragmented scaffolds that are not placed in their respective chromosomes (Kim et al., 2022; Vara et al., 2021). In this respect, we were unable to confidently place 14-32% of sequences in each genome. Despite this, these assemblies are still of high enough quality to contain the expected number of chromosomes in all three species, with an N50 range of 43-53Mbp.

We used the genome assemblies to design 4 chromosome-specific probes (Ch01A, Ch01B, Ch05, and Ch20). Their application to metaphase spreads of the three species confirmed some features of the assemblies, most notably the fusion of two chromosomes (Ch01A and Ch01B) into a single chromosome (Ch01) in *C. taenia*. Interestingly, however, while both Ch01A, Ch01B appear as small chromosomes in *C. elongatoides* and *C. tanaitica*, they appear to not be acrocentric, with a break in the signal of the probes suggesting the existence of a small pericentromeric region in both chromosomes. This in turn suggests that their fusion into Ch01 of *C. taenia* involved not only merging, but also some fine-scale rearrangement. This is in line with suggested inversions and translocations along both Ch01 - Ch01A & Ch01B (see and compare Figure 6 and Figure 3A, 3B).

Our comparison to published chromosome-level assemblies suggests that the syntenic groups of the Cobitoidea suborder appear highly conserved, with most previously studied species demonstrating 25 elements in haploid state and very few interchromosomal rearrangements. However, fusions of two elements have been identified in the genome assemblies of *Paramisgurnus dabryanus* (2n=48, L. Zhang et al., 2025) and *Oreonectes platycephalus* (2n=48, Wang et al., 2025). A closer investigation of these fused chromosomes reveals their independent origin from the fusion observed in *C. taenia* (Supplementary Table S12).

While the number of chromosome-level genome assemblies from non-model species has grown exponentially, it remains taxonomically and geographically biased. This bias is evident in Cypriniformes, a diverse and economically significant order of Old-World freshwater fishes. As of March 2025, the NCBI Genome database lists 306 genomes of Cypriniformes, including 89 genomes from species of the family Cyprinidae (which has around 1,780 species in total) and only 5 species from the family Cobitidae (which has approximately 260 species). This is even though the Cobitidae family, which diverged from its common ancestor ∼20–30 Mya (Šlechtová et al., 2021), spans the entire Palearctic region. This bias likely stems from the economic importance of the cyprinids, particularly carps, which are farmed extensively worldwide. Loaches, on the other hand, are generally non-commercial, with exceptions in China and Japan, explaining why the available genomes primarily originate from East Asian species.

Such a taxonomic and geographic imbalance is understandable given the costs and labor required for high-quality chromosome-level assemblies. However, it significantly limits large-scale comparative genomic studies by underrepresenting or omitting entire deeply diverged clades. In this context, the present study, delivering chromosome-level genomes of *Cobitis taenia*, *C. tanaitica*, and *C. elongatoides*, more than doubles the number of available Cobitidae genomes. Crucially, it addresses a key taxonomic and geographic gap by providing genomes from the Western Cobitidae lineage (Perdices et al., 2016), which diverged from the nearest available Asian *Misgurnini* species during the Oligocene epoch.

Beyond merely filling this gap, our assemblies—supported by additional work on Pool-Seq and chromosome-specific probe design—have leveraged the unique potential of this dataset to advance our understanding of the intricate links between speciation, hybridization, asexuality, and hybrid sterility at two distinct levels. First, they enabled the detection and characterization of sex chromosomes in these closely related but reproductively divergent hybridizing species. Second, they facilitated a deeper understanding of the mechanisms underpinning chromosomal incompatibilities in hybrids.

### Structural variant evolution

Inversions are structural mutations well known for suppressing recombination in heterozygous states, thereby creating barriers to genetic exchange between chromosomes within populations. A striking feature of *Cobitis* genome evolution is the prevalence of intrachromosomal structural rearrangements over interchromosomal changes, including numerous pericentric and paracentric inversions. Even between the recently diverged species *C. taenia* and *C. tanaitica* (0.5–1.5 Mya), we identified over 50 large (over 1 Mb) inversions (Figure 3A, Supplementary Tables S7, S8), including pericentric ones on Ch01a and Ch01b, confirmed by changes in centromere position from the cytogenetic methods (Figure 6).

Surprisingly, the number of large interchromosomal translocations remains low, limited to a single Ch01a/01b fusion, apomorphic to *C. taenia* as discussed above. The distribution of inversions appears non-random across chromosomes: while some chromosomes (e.g., Ch15, Ch16, Ch21, Ch24) maintain relatively conserved synteny, others (e.g., Ch01b, Ch4, Ch10) exhibit extensive reshuffling (Figure 3; Supplementary Figure S1). This chromosomal reshuffling may influence meiotic pairing frequencies and bivalent formation in hybrids.

Recent studies have demonstrated that inversions tend to fix more frequently in population zones where reproductively isolated species overlap (Hooper & Price, 2017). In such cases, inversions may be positively selected as they preserve loci’s synteny by suppressing recombination in hybrids. Our observations align with this hypothesis. Future pan-genomic projects will target the distribution and population structure of inversions within *Cobitis* species and explore their role in gonosome divergence.

### Repeatome evolution

As in other cypriniform fishes (Shao et al., 2019), *Cobitis* appears to have a relatively high transposable element (TE) content, which likely explains the large genome size in this group. Representatives of both Class I and Class II TEs are almost equally dominant and continue to expand. Similar to other fish genomes, *Cobitis* species are poor in SINEs (Sotero-Caio et al., 2017). However, like cyprinids (Shao et al., 2019), *Cobitis* exhibits a high level of TE diversity.

The *Cobitis* genome has a relatively high LTR content (17-19%) compared to other teleosts, including *Danio rerio* (5%) (Shao et al., 2019) and *Paramisgunus dabryanus* (7.5%) (Zhang et al., 2025). Notably, *C. taenia* shows evidence of a recent transpositional burst involving DNA transposons (hAT), LTR retroelements (Gypsy), and LINE retrotransposons (L2). Compared to many other teleosts, *Cobitis* appears to be undergoing an ongoing transposon expansion, where the rate of active transposon accumulation surpasses the rate of their decline. Even in *C. elongatoides*, with a recent decline in activity, this is still true.

Bursts of TEs have been suggested to play an important role in population diversification and speciation (Jurka et al., 2011; Oliver & Greene, 2011; Platt et al., 2014), and the high TE activity observed in *Cobitis* aligns well with this hypothesis, assuming a contribution of TE to high number of genomic structural variants, rapid karyotype evolution as well as speciation in this group.

### Hybridizing species have nonhomologous genetic sex determination systems

It has also been recognised that mechanisms of sex determination, while being of fundamental importance for species, evolves far more dynamically than previously believed (Heule et al., 2014; Mank & Avise, 2009). The currently available TreeOfSex database v.1 lists 40 cypriniformes species for which sex determination has been investigated. Of these only three previous papers suggested the existence of genetic sex determination (GSD) in 4 species using purely cytogenetic methods throughout the entire Cobitoidei suborder, namely X0 (Vasil’eva & Vasil’ev, 1998), ZW (Sharma & Tripathi, 1988) and multiple sex chromosome X1X2Y (Saitoh, 1989). Additional two recent papers (Wang et al., 2025; L. Zhang et al., 2025) indicated presence of sex chromosomes in two Asian loaches, the XY system in *Oreonectes platycephalus* and ZW system in *Paramisgurnus*, respectively.

Our study brings robust evidence for the presence of GSD in this group of fish, clearly identifying candidate genomic regions/chromosomes responsible for this in each species and validating these predictions with PCR primers in two of the species, which enables further comparative studies within this dynamically evolving field of research. Using Pool-Seq of dozens of male and female individuals in two species and individual male and female sequencing for the third species our study revealed dynamic evolution of GSD within *Cobitis sensu stricto*, namely no signal (*C. taenia*) or ambiguous signal (*C. tanaitica*) on two independent linkage groups, while relatively well differentiated XY chromosomes in Ch01A’s linkage group of *C. elongatoides*, which contains a gene previously associated with the GSD in the blue tilapia and four additional genes regulating the sex determination cascade in vertebrates (Supplementary Table S4). This suggests a rapid turnover of sex chromosomes within the lineage which diverged less than 10 Mya (divergence of *C. elongatoides* from the other species) or even more recently, if we consider that *C. taenia* and C. *tanaitica* diverged in the last 2 Mya (Janko et al., 2018) and appear to have different sex chromosomes (Ch05 and Ch06 respectively).

The observed turnover of sex chromosomes among closely related *Cobitis* species aligns with the well-documented evolutionary plasticity of GSD systems in teleost fishes (Heule et al., 2014; Mank & Avise, 2009). Moreover, the sex chromosomes identified in other cobitoid species (XY in *Oreonectes platycephalus* and ZW in *Paramisgurnus dabryanus*) originated from different syntenic groups (Supplementary Table S12).

However, our finding that *C. elongatoides* utilizes a GSD system (XY on Ch01A) that is non-homologous to the putative sex-determining regions in *C. tanaitica* (XY on Ch06) and *C. taenia* (XY on Ch05) adds a significant dimension to understanding the inherent link between interspecific hybridization and the evolution of asexuality and polyploidy as hybridization between *C. elongatoides* and the other two species consistently produces sterile males and fertile, clonally reproducing females. Such asymmetric patterns are not unique to *Cobitis* but rather have been consistently reported across diverse vertebrate taxa following hybridization events and have long suggested a potential role for sex chromosomes in linking hybrid sterility and asexuality (Stöck et al., 2021). However, in most studied systems involving asexual hybrids, sex chromosomes remained undifferentiated, unknown, or poorly characterized, leaving this hypothesis largely speculative until now (Stöck et al., 2021).

While such asymmetries superficially align with Haldane’s Rule, especially given current evidence for male heterogamety in loaches, the fertility of hybrid females is not maintained through typical meiotic repair mechanisms but arises via PMER — a cellular mechanism where germ cells duplicate their chromosomes before meiosis, allowing bivalents to form between identical chromosomal copies. This bypasses meiotic pairing issues and restores fertility. Crucially, however, PMER is restricted to female hybrids, which subsequently reproduce clonally, while hybrid males remain sterile. Notably, transplantation experiments have demonstrated that PMER can be reactivated in spermatogonial cells of hybrid males if these cells develop in a female gonadal environment and transdifferentiate into oogonia that may subsequently give rise to unreduced oocytes (Tichopád et al., 2022). While such a finding leads to persuasion that the ability to initiate asexual reproduction via PMER is primarily dictated by tissue-specific cues from the female gonadal environment, our discovery adds a new layer of complexity to this. It demonstrates that GSD systems in hybrids combine fundamentally different systems among parental species. Such an interplay between GSD and the female gonadal environment hints at a deeper integration between genetic triggers, epigenetic regulation, and cellular signalling pathways in enabling PMER. It opens a hypothesis to test that while the female-specific gonadal environment acts as a permissive factor for PMER, the genetic sex determination system may serve as an upstream regulatory mechanism, influencing how these cellular pathways are executed in hybrids. Our findings, therefore, highlight an exciting new research avenue into the genetic and molecular basis of PMER and the broader evolutionary consequences of sex chromosome turnover in hybrid systems.

### Chromosome Pairing in Hybrids: Insights into Hybrid Sterility and Asexuality

The assembly of chromosome linkage groups and the design of chromosome-specific probes allowed us to investigate meiotic chromosome pairing in hybrid males, providing unprecedented insights into the chromosomal basis of hybrid sterility and asexual reproduction. Such analyses, examining the pairing rates of individual chromosomes in hybrid meiosis, remain technologically challenging and are rarely performed. Yet, they address a fundamental aspect of hybrid sterility models, which suggest that individual chromosomes vary in their contributions to meiotic success or failure (Bhattacharyya et al., 2013; Forejt & Jansa, 2023). Our findings align with this hypothesis, showing that orthologous chromosome pairs (Ch01A, Ch01B, Ch05, and Ch20) exhibit significantly different rates of bivalent formation in hybrid males (Figure 7D). For example, while two-thirds of spermatocytes contained a bivalent between *C. elongatoides*-derived Ch01B and its ortholog from *C. taenia*, only 2 out of > 300 cells displayed pairing between the Y-linked Ch01A from *C. elongatoides* and its ortholog from *C. taenia*. Analogously, for Ch05 and Ch20, given an average of five bivalents per spermatocyte (Dedukh et al., 2020) and assuming random pairing, approximately 25 cells out of 120 and 117 inspected, respectively, would be expected to contain bivalents of these chromosomes. However, far fewer such cells were observed, indicating systematic biases in pairing success. In addition, significant interindividual differences were found between both hybrid males analyzed in terms of pairing success of at least one chromosome (Ch01B), suggesting that variability in pairing dynamics might reflect unique characteristics of this chromosome or its interaction with individual-specific factors.

These results, the first of their kind in an asexually reproducing vertebrate, gain additional importance when interpreted within the context of asexual gametogenesis pathways. Studies have shown that, even in evolutionarily successful clones, only a minority of gonial cells undergo PMER, while most hybrid female oogonia and all male spermatogonia fail to pair orthologous chromosomes and stall at the first meiotic checkpoint (Dedukh et al., 2020, 2021). Despite this widespread failure, each gonocyte still contains several fully formed bivalents, with their numbers varying between individuals and sexes (∼5 bivalents in hybrid males and ∼16 in hybrid females). This variability raises critical questions about whether specific chromosomes systematically contribute more or less to pairing success and why such differences persist between males and females.

Furthermore, accumulating evidence suggests that asexual hybrids accumulate a loss of heterozygosity (Janko et al., 2021; Jaron et al., 2021; Tucker et al., 2013; Warren et al., 2018). Our results propose an exciting hypothesis: if individual chromosomes differ in their likelihood to form orthologous bivalents during PMER, certain linkage groups may be disproportionately affected by gene conversion events. This non-random pairing propensity could predict non-random distributions of loss of heterozygosity across the genomes of asexual hybrids.

### Conclusion

Our study highlights how integrating chromosome-level genome assemblies, molecular cytogenetics, and meiotic analysis can illuminate the mechanisms underlying hybrid sterility and asexuality. We show that hybridizing *Cobitis* species differ in their sex determination systems, accumulate extensive structural variants, and exhibit nonrandom, chromosome-specific pairing affinities during male meiosis. Notably, the frequent mispairing of the Y-derived Ch01A points to structural or regulatory incompatibilities as potential barriers to normal gametogenesis. Together, these findings underscore how genome divergence between parental species may shape reproductive outcomes in hybrids and pave the way for future research into the chromosomal basis of asexuality.

## Acknowledgements

Authors are profoundly obliged to our greatest technicians, Š. Pelikanová, J. Machová and P. Šejnohová. The study was supported by the Czech Science Foundation Project No. 24-12217S. Institute of Animal Physiology and Genetics receives support from Institutional Research Concept, Grant/ Award Number: RVO67985904. ARH is founded by the Spanish Ministry of Science and Innovation (PID2020-112557GB-I00 funded by AEI/10.13039/501100011033), the Agència de Gestió d’Ajuts Universitaris i de Recerca, AGAUR (2021SGR00122) and the Catalan Institution for Research and Advanced Studies (ICREA). L.A-G. and G.P. were supported by FPI predoctoral fellowships from the Ministry of Economy, Industry, and Competitiveness (PRE-2018-083257 and PRE-C-2021-0083, respectively). ZH was supported by the Grant Agency of Charles University (grant number 314222) and also SVV 260684/2024.

## Notes

### Competing Interest Statement

The authors have declared no competing interest.

