## Supplementary Figures S1-S3 for "Sex Chromosome Turnover and Structural Interspecific Genome Divergence Shapes Meiotic Outcomes in Hybridizing *Cobitis*"

### Sex Chromosome Turnover and Structural Interspecific Genome Divergence Shapes Meiotic Outcomes in Hybridizing Cobitis

S. A. Schlebusch, V. Trifonov, Z. Halenková, M. Klianitskaya, D. Dedukh, A. Ruiz Herrera, L. Álvarez González, G. Pujol Infantes, E. Hřibová, L. Andjel, O. Bartoš, P. Pajer, T. Tichopád, D. Kulik, J. Kotusz, M. Kaštánková Doležálková, A. Bohne, A. Marta, P. Horna, R. Reifová, Y. Guiguen, J. Pačes, K. Janko

**Supplementary Figure S1**: Dotplot analysis of genomic homologies between *Cobitis elongatoides*, *C. taenia*, and *C. tanaitica.* Dotplots depict pairwise genomic comparisons between *C. elongatoides* (E), *C. taenia* (T), and *C. tanaitica* (N), illustrating sequence homology and structural variation. Each panel represents a pairwise alignment: (A) *C. elongatoides* vs. *C. taenia*, (B) *C. taenia* vs. *C. tanaitica* and (C) *C. elongatoides* vs. *C. tanaitica*. Diagonal lines indicate regions of synteny, while disruptions or scattered points suggest structural rearrangements such as inversions, translocations, or duplications. The density and continuity of dot patterns reflect the level of sequence similarity and collinearity between species.

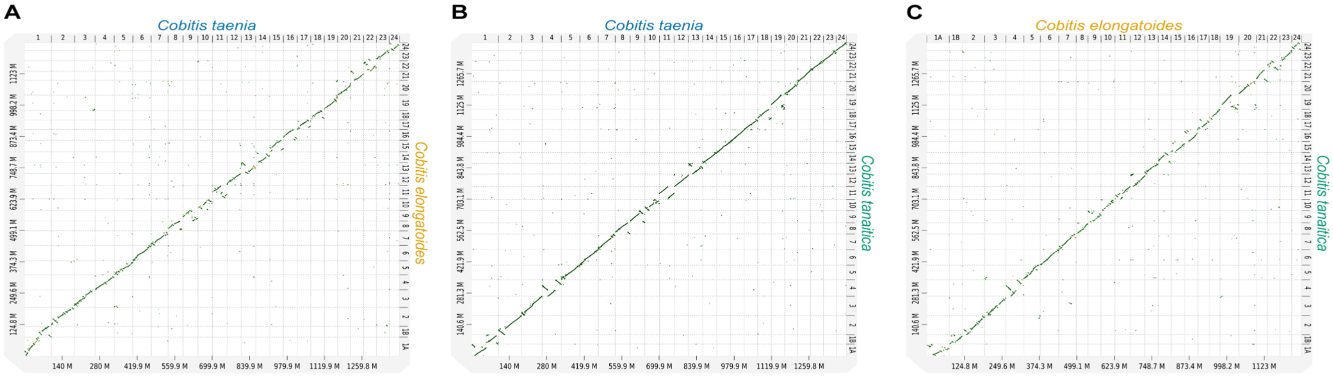
A
Cobitis taenia
Cobitis elongatoides
140 M
280 M
419.9 M
559.9 M
699.9 M
839.9 M
979.9 M
1119.9 M
1259.8 M
B
Cobitis taenia
Cobitis tanaitica
140 M
280 M
419.9 M
559.9 M
699.9 M
839.9 M
979.9 M
1119.9 M
1259.8 M
C
Cobitis elongatoides
Cobitis tanaitica
124.8 M
249.6 M
374.3 M
499.1 M
623.9 M
748.7 M
873.4 M
998.2 M
1123 M

**Supplementary Figure S2**: Genome synteny plot. Syntenic blocks on homologous chromosomes are depicted in light grey while interchromosomal blocks are highlighted in pink.

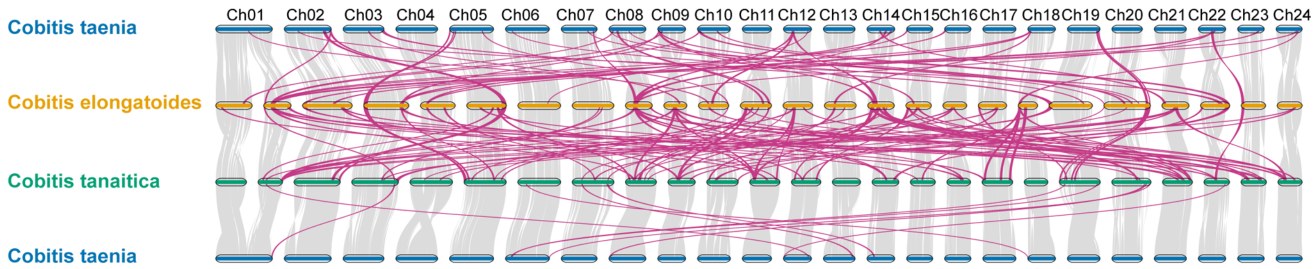

Cobitis taenia
Cobitis elongatoides
Cobitis tanaitica
Cobitis taenia
Ch01
Ch02
Ch03
Ch04
Ch05
Ch06
Ch07
Ch08
Ch09
Ch10
Ch11
Ch12
Ch13
Ch14
Ch15
Ch16
Ch17
Ch18
Ch19
Ch20
Ch21
Ch22
Ch23
Ch24

**Supplementary Figure S3**: Chromosome painting of Ch05 and Ch20 (A-C) as well as Ch01A and Ch01B (D-E) on meiotic metaphases of *C. taenia* (A, D), *C. tanaitica* (B, E), *C. elongatoides* (C, F). Ch05 and Ch20 (a-c) indicate two bivalents in all species. Chromosome painting of Ch01A and Ch01B indicate one bivalent in *C. taenia* (D) while two bivalents in *C. tanaitica* (E) and *C. elongatoides* (F). Chromosome are stained by DAPI (blue). Scale bar = 10 μl.

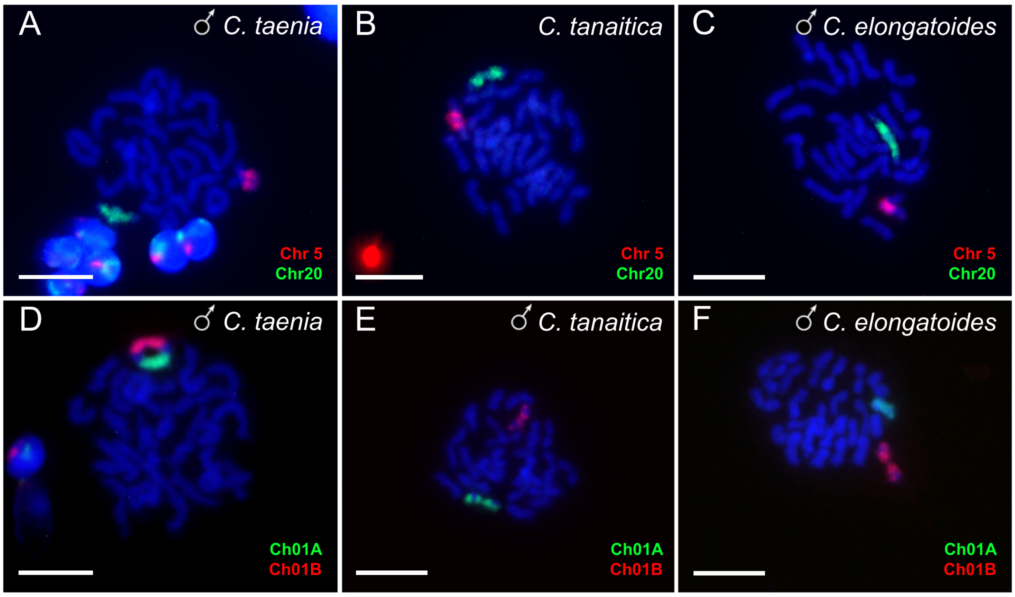

A
♂ C. taenia
Chr 5
Chr20
B
C. tanaitica
Chr 5
Chr20
C
♂ C. elongatoides
Chr 5
Chr20
D
♂ C. taenia
Ch01A
Ch01B
E
♂ C. tanaitica
Ch01A
Ch01B
F
♂ C. elongatoides
Ch01A
Ch01B
